## Supplementary Materials for "Thermal Contrast Enhancement Predicts Paradoxical Heat Sensation"

**Temporal Contrast Enhancement in Thermosensation: A Framework for Understanding Paradoxical Heat Sensation**

**Supplementary Materials**

***Supplementary Data:* Distribution of TSL thresholds and TCF for both innocuous and noxious tasks**

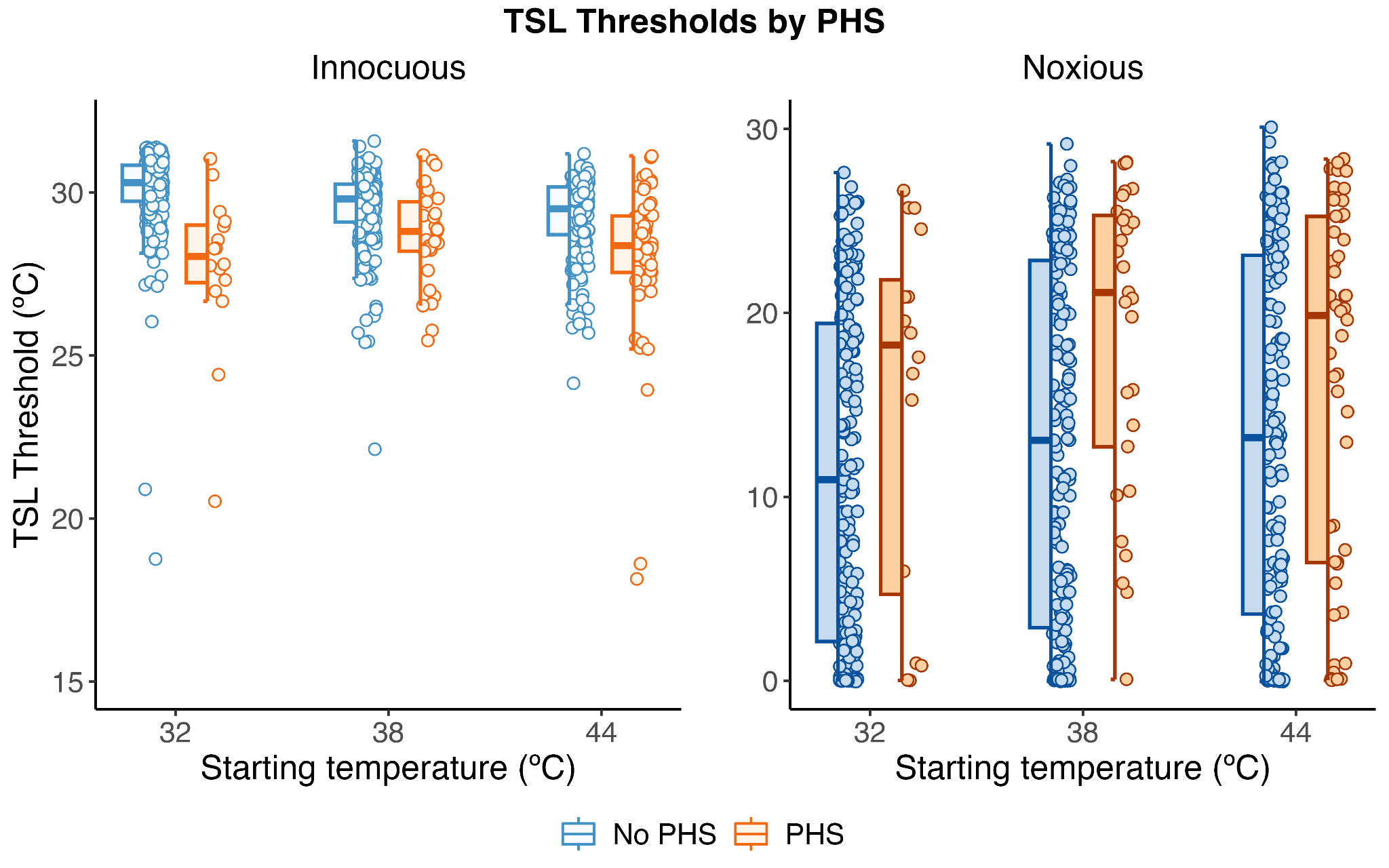

***Supplementary Figure 1****: Innocuous and noxious TSL threshold values for each contrast condition (starting temperature) for veridical experience of cold and PHS.*

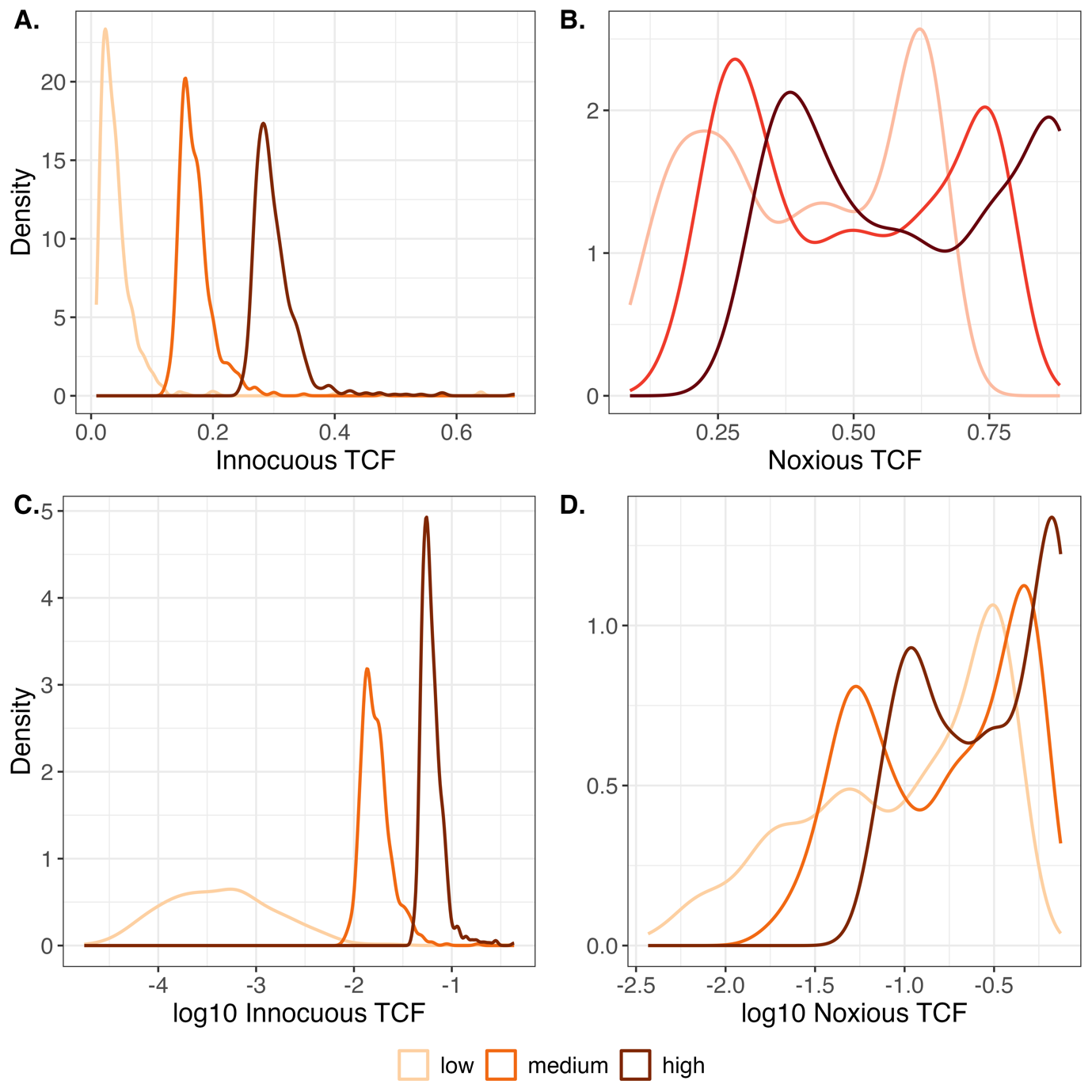

***Supplementary Figure 2****: Density plots of the distribution of (A) innocuous and (B) noxious TCF and changes in distribution by calculating the log10 of (C) innocuous and (D) noxious TCF, which was used in the final logistic regression. As the TCF is a standardised threshold value, the distribution of innocuous and noxious TSL thresholds match that of the raw TCF, just over a different scale.*

***Supplementary Note 1:* Full results for fixed effects in all models presented in the manuscript with associated omnibus tests**

Model 1A: PHS ~ contrastCondition * task + (1|ID)

| *1A Regression Parameters* | | | | | |
| --- | --- | --- | --- | --- | --- |
|  | **β** | **SE** | **z-value** | ***p*-value** | **OR [95% CI]** |
| **Intercept** | -4.29 | .29 | -14.86 | <.001 | .01 [.01 - .02] |
| **32 vs. 38ºC** | .57 | .29 | 1.97 | .05 | 1.77 [1.00 – 3.14] |
| **32 vs. 44ºC** | 1.62 | .26 | 6.12 | <.001 | 5.05 [3.00 – 3.14] |
| **Innoc. vs nox.** | .90 | .28 | 3.23 | .001 | 2.46 [1.43 – 4.27] |
| **38ºC * nox.** | -.94 | .38 | -2.45 | .01 | .39 [.19 - .83] |
| **44ºC * nox.** | -2.38 | .38 | -6.26 | <.001 | .09 [.04 - .20] |

| *1A Omnibus Test* | | | |
| --- | --- | --- | --- |
|  | **χ^2^** | **df** | ***p*-value** |
| **contrastCondition** | 12.52 | 2 | .002 |
| **task** | 3.24 | 1 | .07 |
| **contrastCondition*task** | 40.47 | 2 | <.001 |

Model 2A: innocuousTSL ~ contrastCondition + trial_z + (1|ID)

| *2A Regression Parameters* | | | | | |
| --- | --- | --- | --- | --- | --- |
|  | **β** | **SE** | **df** | **t-value** | ***p*-value** |
| **Intercept** | 29.85 | .10 | 438.12 | 307.28 | <.001 |
| **32 vs. 38ºC** | -.46 | .10 | 1661.00 | -4.86 | <.001 |
| **32 vs. 44ºC** | -.95 | .10 | 1661.00 | -9.95 | <.001 |
| **Trial** | -.16 | .04 | 1661.28 | -4.19 | <.001 |

| *2A Omnibus Test* | | | |
| --- | --- | --- | --- |
|  | **χ^2^** | **df** | ***p*-value** |
| **contrastCondition** | 98.94 | 2 | <.001 |
| **trial_z** | 17.58 | 1 | <.001 |

Model 2B: noxiousTSL ~ contrastCondition + trial_z + (1|ID)

| *2B Regression Parameters* | | | | | |
| --- | --- | --- | --- | --- | --- |
|  | **β** | **SE** | **df** | **t-value** | ***p*-value** |
| **Intercept** | 11.72 | .64 | 221.76 | 18.26 | <.001 |
| **32 vs. 38ºC** | 1.86 | .20 | 1661.00 | 9.05 | <.001 |
| **32 vs. 44ºC** | 2.12 | .20 | 1661.00 | 10.36 | <.001 |
| **Trial** | -.92 | .08 | 1661.05 | -11.04 | <.001 |

| *2B Omnibus Test* | | | |
| --- | --- | --- | --- |
|  | **χ^2^** | **df** | ***p*-value** |
| **contrastCondition** | 127.33 | 2 | <.001 |
| **trial_z** | 121.89 | 1 | <.001 |

Model 2C: innocuousPHS ~ innocuousTSL * noxiousTSL + (1|ID)

| *2C Regression Parameters* | | | | | |
| --- | --- | --- | --- | --- | --- |
|  | **β** | **SE** | **z-value** | ***p*-value** | **OR [95% CI]** |
| **Intercept** | -7.56 | .70 | -11.30 | <.001 | <.01 [<.01 - <.01] |
| **Innocuous TSL** | -1.56 | .11 | -14.18 | <.001 | .21 [.17 – .26] |
| **Noxious TSL** | .94 | .22 | 4.37 | <.001 | 2.56 [1.68 – 3.91] |
| **Innoc. * nox.** | -.68 | .10 | -7.02 | <.001 | .51 [.42 – .61] |

Model 3A: innocuousPHS ~ contrastCondition + (1|ID)

| *3A Regression Parameters* | | | | | |
| --- | --- | --- | --- | --- | --- |
|  | **β** | **SE** | **z-value** | ***p*-value** | **OR [95% CI]** |
| **Intercept** | -4.71 | .39 | -12.43 | <.001 | .01 [<.01 - .02] |
| **32 vs. 38ºC** | .61 | .30 | .207 | .04 | 1.83 [1.02 – 3.29] |
| **32 vs. 44ºC** | 1.72 | .28 | 6.19 | <.001 | 5.58 [3.24 – 9.61] |

| *3A Omnibus Test* | | | |
| --- | --- | --- | --- |
|  | **χ^2^** | **df** | ***p*-value** |
| **contrastCondition** | 45.89 | 2 | <.001 |

Model 3B: innocuousPHS ~ log(innocuousTCF) + (1|ID)

| *3B Regression Parameters* | | | | | |
| --- | --- | --- | --- | --- | --- |
|  | **β** | **SE** | **z-value** | ***p*-value** | **OR [95% CI]** |
| **Intercept** | -1.34 | .38 | -3.51 | <.001 | .26 [.12 - .55] |
| **Innocuous TCF** | 1.37 | .19 | 7.24 | <.001 | 3.94 [2.72 – 5.70] |

Model 3C: innocuousPHS ~ log(noxiousTCF) + (1|ID)

| *3C Regression Parameters* | | | | | |
| --- | --- | --- | --- | --- | --- |
|  | **β** | **SE** | **z-value** | ***p*-value** | **OR [95% CI]** |
| **Intercept** | -3.38 | .32 | -10.65 | <.001 | .03 [.02 - .06] |
| **Noxious TCF** | .38 | .30 | 1.26 | .21 | 1.46 [.81 – 2.65] |

Model 3D: innocuousPHS ~ log(innocuousTCF) * log(noxiousTCF) + (1|ID)

| *3D Regression Parameters* | | | | | |
| --- | --- | --- | --- | --- | --- |
|  | **β** | **SE** | **z-value** | ***p*-value** | **OR [95% CI]** |
| **Intercept** | -.91 | .74 | -1.23 | .22 | .40 [.09 – 1.72] |
| **Innocuous TCF** | 2.60 | .49 | 5.35 | <.001 | 13.48 [5.20 – 34.99] |
| **Noxious TCF** | -1.22 | .69 | -1.77 | .07 | .29 [.08 – 1.14] |
| **Innoc. * Nox.** | .46 | .32 | 1.46 | .15 | 1.58 [.85 – 2.94] |

***Supplementary Note 2***

***Supplementary Table 1:*** *Mean QST detection and pain thresholds (ºC) for individuals without PHS compared to those with PHS. No significant relationship was observed between PHS prevalence and cold detection (CDT), warm detection (WDT), cold pain (CPT) and heat pain (HPT) thresholds.*

|  | **No PHS** | | **PHS** | |
| --- | --- | --- | --- | --- |
|  | *Mean* | *SD* | *Mean* | *SD* |
| **CDT** | 30.33 | 0.86 | 30.41 | 1.05 |
| **WDT** | 33.86 | 0.61 | 33.98 | 0.86 |
| **CPT** | 14.23 | 9.11 | 17.80 | 9.44 |
| **HPT** | 42.81 | 3.43 | 42.17 | 3.65 |

**QST model results**

| innocuousPHS ~ CDT + WDT + CPT + HPT + (1\|ID) | | | | | |
| --- | --- | --- | --- | --- | --- |
|  | **β** | **SE** | **z-value** | ***p*-value** | **OR [95% CI]** |
| **Intercept** | -14.25 | 2.65 | -5.38 | <.001 | <.01 [<.01 - <.01] |
| **CDT** | -.28 | 1.36 | -.21 | .84 | .75 [.05 – 10.76] |
| **WDT** | .33 | 1.44 | .23 | .82 | 1.39 [.08 – 23.38] |
| **CPT** | 1.10 | 2.81 | .39 | .70 | 3.01 [.01 – 74.29] |
| **HPT** | .83 | 2.71 | .31 | .76 | 2.27 [.01 – 46.48] |

***Supplementary Note 3:* Trial number affects TSL thresholds but not PHS**

We conducted an extension of Model 1A in the manuscript with the inclusion of trial number (z-scored) (Model S1). This was to account for the assumption that the probability PHS may be modulated by the trial number. We found no significant effect of trial on PHS (z = -0.67, *p* = .50, OR = 0.92, 95% CI = 0.71 – 1.18) and the addition of trial number did not significantly improve upon Model 1A (*p* = .91).

$PHS \sim contrastCondition*task+trial\_z + (1|ID)$	(Model S1)

In addition to this, we included trial number (z-scored) into Models 2A and 2B. Both innocuous and noxious TSL temperatures decreased with increasing trial number (innocuous TSL: t_438.12/1661.28_ = -4.19, β = -.16, *p* < .001; noxious TSL: t_221.76/1661.05_ = -11.04, β = -.92, *p* < .001).

***Supplementary Note 4:* Effect of age and gender on TSL thresholds and PHS**

We explored the possible effects of age and gender (male or female) on both TSL thresholds and PHS by adding these predictors to models 2A, 2B and 3D in the manuscript (Models S2, S3 & S4). Neither age nor gender significantly affected innocuous (age: β = -.01, t = -.56, df = 213.10/205, *p* = .58; gender: β = -.15, t = -.90, df = 213.10/205, *p* = .37) or noxious (age: β = -.21, t = -1.74, df = 205.60/205, *p* = .08; gender: β = -1.15, t = -.89, df = 205.60/205, *p* = .37) TSL thresholds. PHS probability was also not significantly affected by age (z = -.63, *p* = .53, OR = .98, 95% CI = .91 – 1.05) or gender (z = 1.74, *p* = .08, OR = 1.93, 95% CI = .92 – 4.04).

$innocuousTSL \sim contrastCondition+trial_{z}+age+gender+ (1|ID).$	(Model S2)

$noxiousTSL \sim iontrastCondition+trial_{z}+age+gender+ (1|ID)$	(Model S3)

$PHS \sim innocuousTCF*noxiousTCF+age+gender + (1|ID)$	(Model S4)
